# Methane Cycling Microbes are Important Predictors of Methylmercury Accumulation in Rice Paddies

**DOI:** 10.1101/2025.10.09.681528

**Authors:** Rui Zhang, Alexandre J Poulain, Qiang Pu, Jiang Liu, Mahmoud A. Abdelhafiz, Xinbin Feng, Bo Meng, Daniel S Grégoire

**Affiliations:** Department of Biology, University of Ottawa, Ottawa, Canada; State Key Laboratory of Environmental Geochemistry, Institute of Geochemistry, Chinese Academy of Sciences, Guiyang, China; Department of Chemistry, Carleton University, Ottawa, Canada

**Keywords:** methanogen, methanotroph, mercury, methylmercury, methylation, demethylation, metagenomics

## Abstract

Microbial production of methylmercury from inorganic mercury in rice paddies poses health risks to consumers of this essential dietary staple. Although mercury-methylating communities are well characterized, the microbial guilds contributing to methylmercury accumulation in rice paddies remain unclear. Here, we collected paddy soils across a mercury concentration gradient throughout the rice growing season to identify microbial and environmental factors influencing methylmercury dynamics. We show that *hgcA* gene abundance, the key gene required for methylation, was not a significant predictor of methylmercury concentration in paddy soils. We also show that *merB* gene abundance correlated with methylmercury in mercury-polluted rhizosphere samples. Methane cycling genes were actively expressed, and their beta-diversity was significantly associated with methylmercury levels. Methanogen abundance correlated with higher methylmercury under elevated total mercury concentrations. Analysis of the methanotroph-associated *mbnT* gene, implicated in demethylation, revealed an unexpected positive correlation with methylmercury. Multiple regression and machine learning models converged on mercury bioavailability and methanogen/methanotroph abundances as key predictors of methylmercury, with methanogen-associated *hgcA* gene abundance and methanogen-methanotroph interactions highlighted under flooded, low-redox conditions. These findings suggest that methane-cycling microbes play key roles in methylmercury cycling dynamics and point to management strategies that could simultaneously mitigate mercury pollution and greenhouse gas emissions.

**Importance:** Methylmercury is a microbially-derived neurotoxin that accumulates in rice, which is a global food staple. Predicting mercury’s fate in rice paddies is challenging because of the interplay between microbes responsible for methylmercury cycling and variables that control mercury availability. Our study coupled genomic and geochemical measurements with machine learning to identify the key predictors of methylmercury accumulation in paddy soils. We demonstrate that methanogen and methanotroph abundance, and mercury bioavailability, are major predictors of methylmercury variability in paddies. We show that considering interactions between methane cycling guilds improves our capacity to predict methylmercury accumulation in soils compared to approaches that rely solely on mercury cycling genes. This work can inform remediation strategies for mercury in rice paddies but also wetlands and permafrost where methane and mercury cycling are tightly coupled. Such strategies could provide a solution to simultaneously mitigate methylmercury exposure and reduce greenhouse gas emissions amid global environmental change.

## Introduction

Mercury (Hg) is a metal primarily emitted as gaseous elemental Hg (Hg(0)) from natural and anthropogenic sources and deposits in ecosystems worldwide, where it undergoes oxidation to Hg(II) and subsequent microbial conversion to methylmercury (MeHg), a potent neurotoxin (1). MeHg accumulates in rice, a global food staple, with particularly elevated concentrations in paddies near mining, smelting, and industrial areas with elevated Hg(0) emissions (2). The global atmospheric transportation of Hg could also affect rice paddies away from pollution sources (3). Since paddy soil represents a major source of MeHg in rice grains (4), predicting MeHg accumulation in rice paddies is critical for mitigating Hg exposure through diet.

Net MeHg accumulation in rice paddies represents an interplay between methylation and demethylation processes (5). Hg methylation is facilitated by anaerobic microbes harboring the *hgcAB* gene cluster (6–8), while demethylation occurs via the Hg resistance (*mer*) operon– dependent pathways (e.g., *merB*/*merA*) and *mer*-independent microbial pathways (5). These processes are also connectively influenced by environmental variables such as Hg bioavailability, redox, dissolved organic matter (DOM), sulfide (5,9–12) and other microbial players indirectly (13).

While Hg methylating communities have been extensively characterized in rice paddies and other ecosystems (14–19), MeHg poorly correlates with Hg-cycling genes (e.g., *hgcA*, *merA*, *merB*) and *hgcA* transcripts (20). This disconnect highlights the need for integrated approaches that consider multiple variables simultaneously when predicting what controls MeHg accumulation in rice.

Recent evidence suggests that methane-cycling microbes play underappreciated roles in mediating MeHg dynamics. In rice paddies near Hg-mining areas, methanogens emerged as critical players controlling MeHg levels (14,15,21), potentially contributing to both methylation and demethylation, simultaneously (14,22,23). Furthermore, methanotrophs were experimentally shown to degrade MeHg at picomolar to nanomolar concentrations under near- neutral pH conditions typical of rice paddies, using a methanobactin-mediated mechanism distinct from the conventional *mer*-dependent pathway (24,25). However, the ecosystem-level significance of methanotrophic MeHg demethylation and its interaction with methanogenic processes in determining net MeHg accumulation has not been evaluated in rice paddy systems.

To address these critical gaps, we aim to (i) identify and rank the relative importance of microbial communities and environmental variables relevant to MeHg turnover as predictors of MeHg accumulation patterns in mining-impacted rice paddies; (ii) characterize the abundance and distribution of methane-cycling microorganisms in relation to MeHg concentrations; and (iii) develop predictive models to inform future mechanistic studies on Hg cycling.

By integrating metagenomic, geochemical, and statistical approaches, we show that methanogen and methanotroph abundances are the leading biological predictor, while Hg bioavailability is the primary environmental determinant of MeHg variability. Machine learning analyses further highlighted the importance of methanogen methylator abundance and a methanogen–methanotroph interaction, particularly under flooded conditions where model performance improved. Together, these findings provide environmental-scale data that guide hypothesis development for broader geographical testing, and inform strategies that may jointly mitigate greenhouse gas emissions and Hg exposure.

## Materials and Methods

### Soil collection, geochemical analyses, and sequencing

Rice paddy soil samples were collected from three sites in Guizhou province, Southwest China: Huaxi (HX, background site), Gouxi (GX, artisanal Hg smelting area), and Sikeng (SK, abandoned Hg mining area). Sampling occurred from May 2021 to April 2022, spanning pre- plantation through post-harvest periods. Surface and rhizosphere soils were collected and processed as follows: individual replicate samples were analyzed separately for geochemical parameters, while replicate samples were combined into composite samples for DNA extraction and subsequent metagenomic analyses. Soil geochemical parameters were analyzed, including Hg species (THg by Cold Vapor Atomic Fluorescence Spectrometry (CVAFS), MeHg by gas chromatography-CVAFS following USEPA methods, and bio-available fraction Hg by the Mg(NO₃)₂ extraction method, anions (by ion chromatography) (26–28), elemental sulfur (by high-performance liquid chromatography), dissolved organic carbon, soil organic matter, and redox-sensitive species (S²⁻, Fe²⁺, Fe³⁺). Recovery rates of the standard reference materials for THg and MeHg analyses were provided in Supplementary Data 1. DNA and RNA extractions were performed using the PowerSoil DNA Isolation Kit and the RNeasy PowerSoil Total RNA Kit (QIAGEN) following the manufacturer’s instructions. Both metagenomic and metatranscriptomic sequencing were performed on an Illumina NovaSeq 6000 platform, producing 150 bp paired-end reads. Sequencing was conducted at Azenta (Suzhou, China). The sequencing aimed for a read depth of approximately 15 Gbp per sample (range: 10.9 – 20 Gbp). Quality metrics are provided in Supplementary Data 2. In total, 50 metagenomic and 6 metatranscriptomic samples were obtained. Details on sample collection, geochemical analyses and sequencing are provided in Supplementary Method 1.

### Genome assembly and binning

Metagenomic data were processed using fastp and FastQC for quality control (Supplementary Data 3) (29,30). Individual assemblies were performed with MetaSPAdes, while site-specific co-assemblies used MEGAHIT (31). Contigs>2000 bp were retained and mapped with BWA-MEM to generate coverage data. Key assembly metrics, such as N50, total contigs, length distribution and mapping rates are provided in Supplementary Data 4. Four binning algorithms (MetaBAT 2, MaxBin 2, CONCOCT, and VAMB) were applied (32–35), yielding 5307 initial bins. Bin quality was assessed with CheckM2 (36), and DAS-Tool (37) was used to select the highest quality bins, followed by dereplication with dRep (38) (ANI threshold 98%). We retained 267 metagenome-assembled genomes (MAGs) with>50% completeness and<10% redundancy following manual refinement in Anvi’o (39). Taxonomic classification was performed using GTDB-Tk (40) based on GTDB (release 214) (Supplementary Data 5()) . The co-assembly approach was chosen over individual assembly due to higher MAG yield and greater taxonomic diversity. Details on genome assembly, binning and software versions are provided in Supplementary Method 2.

### Metatranscriptomic sequence processing

The RNA sequences were trimmed and filtered to a minimum mean quality score of 30 using fastp (29). Forward and reverse sequences were overlapped with a minimum and maximum of 10 and 150 bp, respectively, using FLASH (41). Sequences were subsequently sorted into SSU ribosomal RNA (rRNA), LSU rRNA, and non-rRNA using SortMeRNA (42) against the default reference database (smr_v4.3_default_db). The non-rRNA fraction was mapped to the ORFs identified on the contigs from the corresponding sample and linked with featureCounts results to determine the transcript abundance of *mcrA*, *pmoA*, *mmoX*, *hgcA*, and *merB*. Taxon-specific transcript abundance was calculated by dividing the RNA reads mapped to transcripts by the total RNA reads in the sample, normalized to HMM length.

### Functional gene abundance analyses

For microbial abundance estimation of various functional guilds, including methanogens, methanotrophs, Hg-methylating microbes, and demethylating microbes, we employed a gene-centric approach at the contig level using individual assemblies. We selected individual assemblies over co-assembly to preserve sample-specific information and enable more accurate quantification of functional genes within each sample. While co-assembly is valuable for generating high-quality MAGs by aggregating data across samples, it inherently homogenizes information from multiple samples, potentially obscuring subtle variations in gene abundance at the individual sample level.

Open reading frames (ORFs) were identified using Prodigal (43), short reads were mapped using BWA MEM (44), and reads mapped to ORFs were quantified using featureCounts from Subread package (45). Functional genes were retrieved using Hidden Markov Models (HMMs) through hmmsearch of HMMER software (46). Specifically, the McrA HMM (PF02249) was sourced from PFAM, while PmoA and MmoX HMMs were obtained from the GraftM gene packages (https://data.ace.uq.edu.au/public/graftm/7) with an E-value threshold of 1×10⁻⁵. We conducted protein homology searches using BLASTp against the NCBI non-redundant database (as of Dec 5, 2023) to confirm their identity and refine search results. Queries consisted of HMM hits of the corresponding genes, each limited to the single best hit (-max_target_seqs 1) to focus on the most relevant homolog. McrA, PmoA, and MmoX hits were filtered based on sequence titles indicative of their respective functions (e.g., McrA hits containing "methyl coenzyme M reductase"; PmoA and MmoX hits classified as "methane monooxygenase"). Extra precautions were taken with methane-cycling genes, as PmoA sequence is homologous to other alkane monooxygenases, and it shares homology with the AmoA of ammonia oxidizers. The highest similarity to a known methanotrophic isolate was prioritized.

Methanobactin biosynthesis and transport genes from references genomes, MAGs and metagenomes were identified using HMMs listed from a previous study with predefined threshold values (47).

Hg-cycling genes required specialized validation procedures. *hgcAB* genes were identified using Hg-MATE-Db HMMs (ver. 1.01142021) (48) and filtered for conserved motifs [N(V/I)WCA(A/G)GK] and [C(M/I)ECGA] (6). MerB sequences were retrieved using pre-compiled HMMs (49) and validated for characteristic cysteine and aspartic acid signatures (50).

For additional functional guilds, IRB detection employed HMM profiles from FeGenie (51) to identify genes related to iron reduction, with gene-specific domain E-value cutoffs as defined in the database configuration files. Abundance for IRB was calculated by summing coverage counts of all identified genes and normalizing against total coding sequence (CDS) counts in each metagenome. For SRB quantification, we targeted the *dsrA* gene using the TIGR02064.1 HMM profile with a domain E-value cutoff of 392.9.

Taxonomic classification was conducted using MMseqs2 (52) against GTDB (release 214) (53) with lowest common ancestor strategy. To facilitate cross-sample comparison, we converted taxon-specific coverage values to the percentage of reads normalized to HMM lengths. Total reads numbers in each sample were referred to effective library size values calculated using the edgeR package (54). Details on functional gene analyses and software versions are provided in Supplementary Method 3.

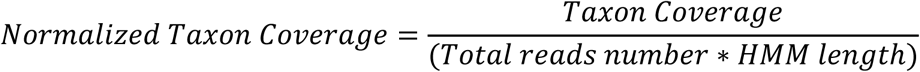

### Complementary metagenomic approaches

Our study employed two complementary metagenomic strategies. Gene-centric analysis quantifies functional potential across all community members, capturing the full breadth of metabolic capacities including genes from low-abundance taxa that may not assemble into complete genomes. Genome-resolved analysis using MAGs provides taxonomic context, linking functional genes to specific organisms and enabling network analyses of population-level responses to environmental changes. Together, these approaches address different scales: community-wide functional potential (gene-centric) and organism-specific contributions (genome-resolved), both essential for understanding Hg cycling processes.

### Statistical analyses

Principal Component Analysis (PCA) of geochemical variables was performed with standardized data, and distance-based redundancy analysis (db-RDA) was used to assess relationships between methane-cycling communities and environmental factors based on Bray-Curtis dissimilarity. Permutational multivariate ANOVA (PERMANOVA) evaluated effects of site and soil compartment on community composition (i.e., beta-diversity), with pairwise comparisons using Bonferroni correction. Multiple linear regression with AICc model selection was employed to evaluate the relative importance of methylation potential (*hgcA* abundance) and Hg bioavailability (F1-Hg) in predicting MeHg concentrations. Seven candidate models representing different hypotheses were tested, and model assumptions were verified through standard diagnostic tests.

MAG relative abundances were calculated by aligning reads to dereplicated genome sets using CoverM (v0.7.0) (55). Time-dependent correlations between MAG abundance and MeHg concentrations were identified using extended Local Similarity Analysis (eLSA) across 11 time points for each site (Supplementary Data 6) (56,57). False Discovery Rate correction was applied using the Benjamini-Hochberg procedure, with significant associations defined as FDR-adjusted p<0.05. Random forest regression models were developed to predict MeHg concentrations from 12 predictor variables (9 microbial functional guilds and 3 environmental parameters) using 100-fold cross-validation with 70:30 train-test splits. Model performance was evaluated using RMSE, MAE, R², and Pearson correlation coefficients. All analyses were conducted in R (v4.3.0) with statistical significance set at α=0.05. Detailed methodological parameters are provided in Supplementary Method 4.

### Code and data availability

The code used for the bioinformatics pipeline is available at https://github.com/rzhan186/gy2021_bioinformatics. Supplementary Methods, Discussions, and Figures are provided in the Supplementary Materials. Supplementary Data and Results can be accessed at https://doi.org/10.6084/m9.figshare.29847869.v3. Raw sequencing data have been deposited in the NCBI Sequence Read Archive under BioProject ID PRJNA1179437.

## Results and Discussion

### Spatial and temporal dynamics of MeHg concentration across the rice paddies

THg and MeHg levels varied significantly across rice paddies (Fig. 1, Fig. S3). Average THg concentrations across the entire sampling period were significantly elevated at the mining- impacted sites SK (42.03±21.5 mg/kg) and GX (21.41±4.04 mg/kg), compared to the background site HX (0.17±0.02 mg/kg). However, average MeHg concentrations over the sampling period were significantly higher at GX (3.44±0.70 μg/kg), followed by SK (2.42±0.76 μg/kg) and HX (0.78±0.21 μg/kg) (Fig. S3). This supports prior studies indicating that THg is a poor predictor of the net MeHg production (58).

**Figure 1.**
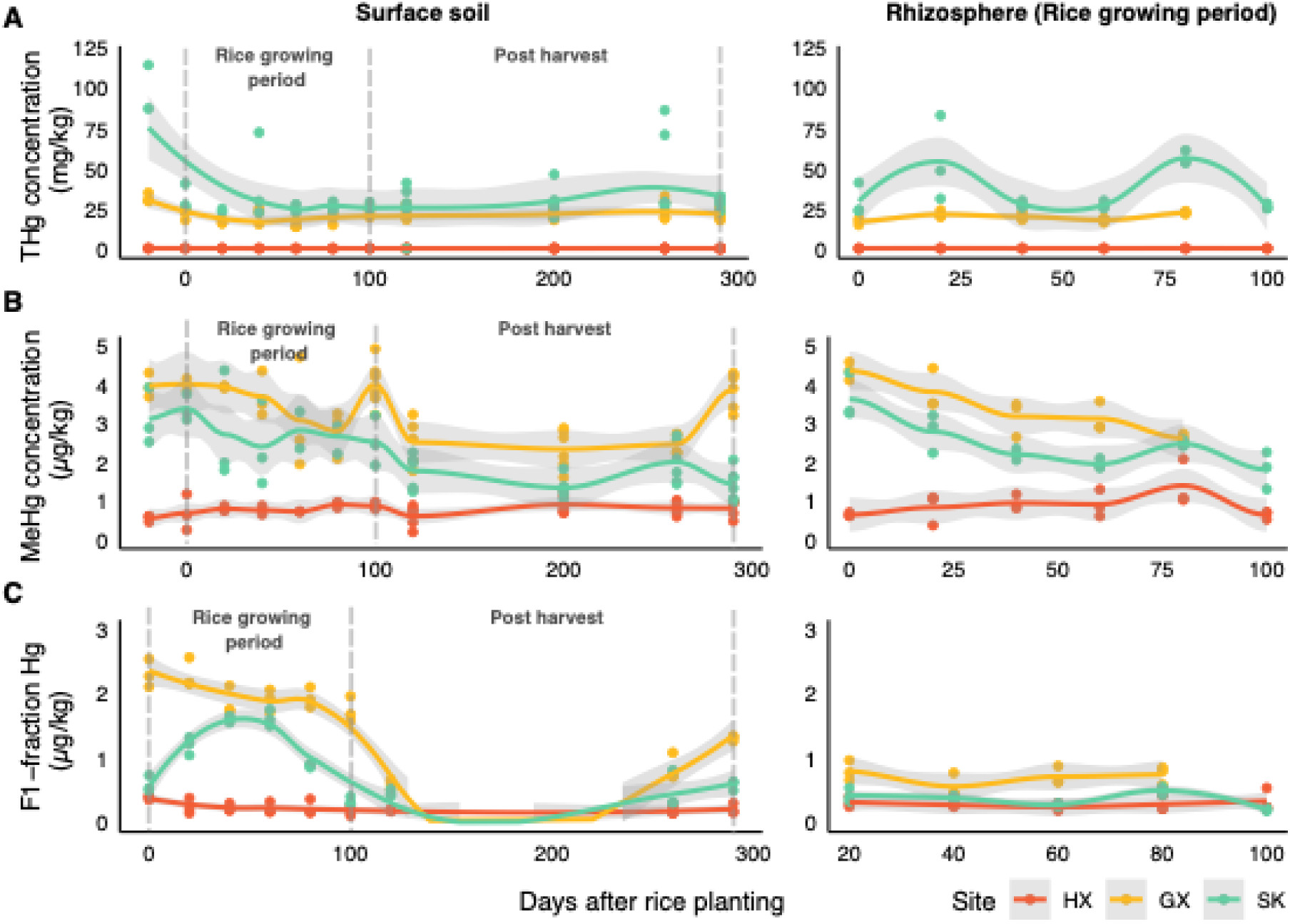
Hg dynamics in sampled rice paddies over the growing season. (A) Total mercury (THg), (B) methylmercury (MeHg), and (C) water-soluble Hg fraction (F1) in surface soils (left panels) and rhizosphere soils (right panels). Data points represent individual measurements from replicate soil samples collected at three sites (HX, GX, and SK), with smoothed trend lines (±95% confidence interval). Note that surface soils were sampled throughout the entire study period (including post-harvest), while rhizosphere soils were only collected during the rice growing season when plants were present, resulting in different x-axis scales. THg is shown in mg/kg, while MeHg and F1-fraction are in μg/kg. Day 0 corresponds to the beginning of the sampling period.

Distinct temporal trends were observed for MeHg at the mining-impacted sites, which exhibited different patterns compared to the relatively stable levels of THg (Fig. 1A). At GX and SK, MeHg peaked at cultivation onset (day 0) and declined throughout the rice-growing season (day 0-80), with further reductions observed post-harvest after drainage. MeHg concentrations were typically higher during the water-saturated season (day 0-100) than in the post-harvest period (day 100-260). A notable increase in MeHg occurred at GX on day 290, coinciding with pre-planting and re-drainage for the next growing season. In contrast, at HX, MeHg levels remained relatively stable overall. Throughout the three sampling sites, no significant differences in MeHg were detected between surface soil and rhizosphere (Mann-Whitney U test, p>0.05). This pattern of fluctuating MeHg in mining-affected sites during cultivation and stability at the background site is previously undocumented, due to the lack of extensive longitudinal sampling. Subsequent sections consider the potential microbial and geochemical drivers contributing to these trends in Hg concentrations.

### Geochemical factors partially explain site-specific MeHg patterns

Geochemical factors were assessed to contextualize their potential impacts on MeHg accumulation (please refer to Fig. S1, Supplementary Result 1 throughout this section). Redox potential measured in overlying water revealed that HX is more oxidizing (109.29±19.12 mV) compared to GX (57.41±37.09 mV) and SK (60.96±20.3 mV). Regardless of soil compartments, DOM concentrations differed significantly between sites, with HX showing the highest levels (0.72±0.31 mg/g). MeHg correlated positively with DOM in surface soil at GX (Spearman’s ρ=0.33) and SK (ρ=0.58), while correlating negatively with DOM SUVA_254_ in SK surface (ρ=-0.56) and rhizosphere (ρ=-0.60) (Fig. S2). The bioavailable fraction of Hg (F1-Hg) exhibited temporal fluctuations that partially paralleled MeHg changes, primarily in GX surface soil (ρ=0.54, p=0.0034, n=27) and SK rhizosphere (ρ=0.62, p=0.014, n=15) (Fig. 1C, Fig. S2). Noticeably, GX initially showed higher F1-Hg than SK, but these differences diminished over the growing season with both sites reaching comparable levels after day 100 (Fig. 1C). This decline aligns with the post-harvest period when rice fields are dried and aerated, promoting immobilization of dissolved Hg species through redox-driven precipitation and adsorption processes (e.g., Fe-oxyhydroxide precipitation and adsorption) (12,59).

Several key factors showed no significant differences between sites. Soil pH displayed similar temporal trends at HX and GX, while SK showed more apparent fluctuation, but with no significant site differences overall (Kruskal-Wallis test, p>0.05, Fig. S3). Terminal electron acceptors including sulfate (avg. ∼100 mg/kg or 100 μM) and nitrate (avg. ∼ 4 mg/kg or 6.45 μM) demonstrated similar temporal patterns across all sites. Hydrogen sulfide remained consistently low (0-5 μM) with no significant site differences (Kruskal-Wallis test, p>0.05, Fig. S3). This suggests similar rates of microbial sulfate reduction and constraints on Hg methylation or bioavailability as sulfide can readily react with Hg(II), forming HgS (60,61). Fe²⁺/total Fe ratios showed similar temporal trends except in HX subsurface porewater, which displayed lower ratios suggesting reduced iron reduction at the depth of rice roots.

The three sites differ primarily in bioavailable Hg content (*i.e.,* F1-Hg) and DOM, which are both variable known to control Hg methylation (62–64). Since DOM affects Hg(II) bioavailability at low sulfide conditions (≲30 μM) (65,66), the consistently low sulfide levels and differences in F1-Hg suggest DOM potentially contributes to observed MeHg fluctuations. Interestingly, DOM aromaticity (i.e., SUVA_254_) showed a pronounced effect at SK, where more lower DOM SUVA_254_, indicating higher DOM bioavailability (67), correlated with higher MeHg in both soil compartments, consistent with previous findings in rice paddy environments (62).

Taken together, our geochemical data suggest that iron and sulfur cycling processes have limited influence on net MeHg accumulation and are insufficient to explain the observed spatiotemporal patterns of MeHg fluctuation across our study sites. While ORP measurements indicate that HX conditions favor microaerophilic to anaerobic processes (denitrification, manganese reduction), and GX and SK are more conducive to iron reduction and sulfate reduction, these differences in redox environments do not adequately account for the MeHg distribution patterns observed. Rather, our findings suggest that net MeHg accumulation is likely controlled by methylating microbes beyond SRB and IRB and active demethylation processes. These results highlight the need to assess the potential contributions of alternative microbial methylators and demethylators to MeHg dynamics in rice paddy environments under the geochemical conditions observed in this study.

### MeHg levels show limited association with *hgcA* gene abundance across metabolic guilds

As the key marker gene of Hg methylation, we searched for and classified *hgcA* in the assembled metagenomes (Fig. 2) to assess methylation potential and identify the microbial players involved. Total *hgcA* abundance was, on average, higher at mining-impacted sites GX (mean=8.4×10⁻⁸±3.0×10⁻⁸; n=16) and SK (mean=8.4×10⁻⁸±2.7 × 10⁻⁸; n=17) compared to the background site (mean=5.5 × 10⁻⁸±1.8 × 10⁻⁸; n=16), aligning with the higher MeHg levels observed at contaminated sites (Fig.1, Fig. 2). Metatranscriptomic analysis confirmed active *hgcA* expression across all sites (n=6), with similar expression in surface soils (∼1.0×10⁻⁸) but substantially higher expression in GX and SK rhizospheres (∼1.4–1.7×10⁻⁸), supporting the functional relevance of the metagenomic potential (Fig. S4A).

**Figure 2.**
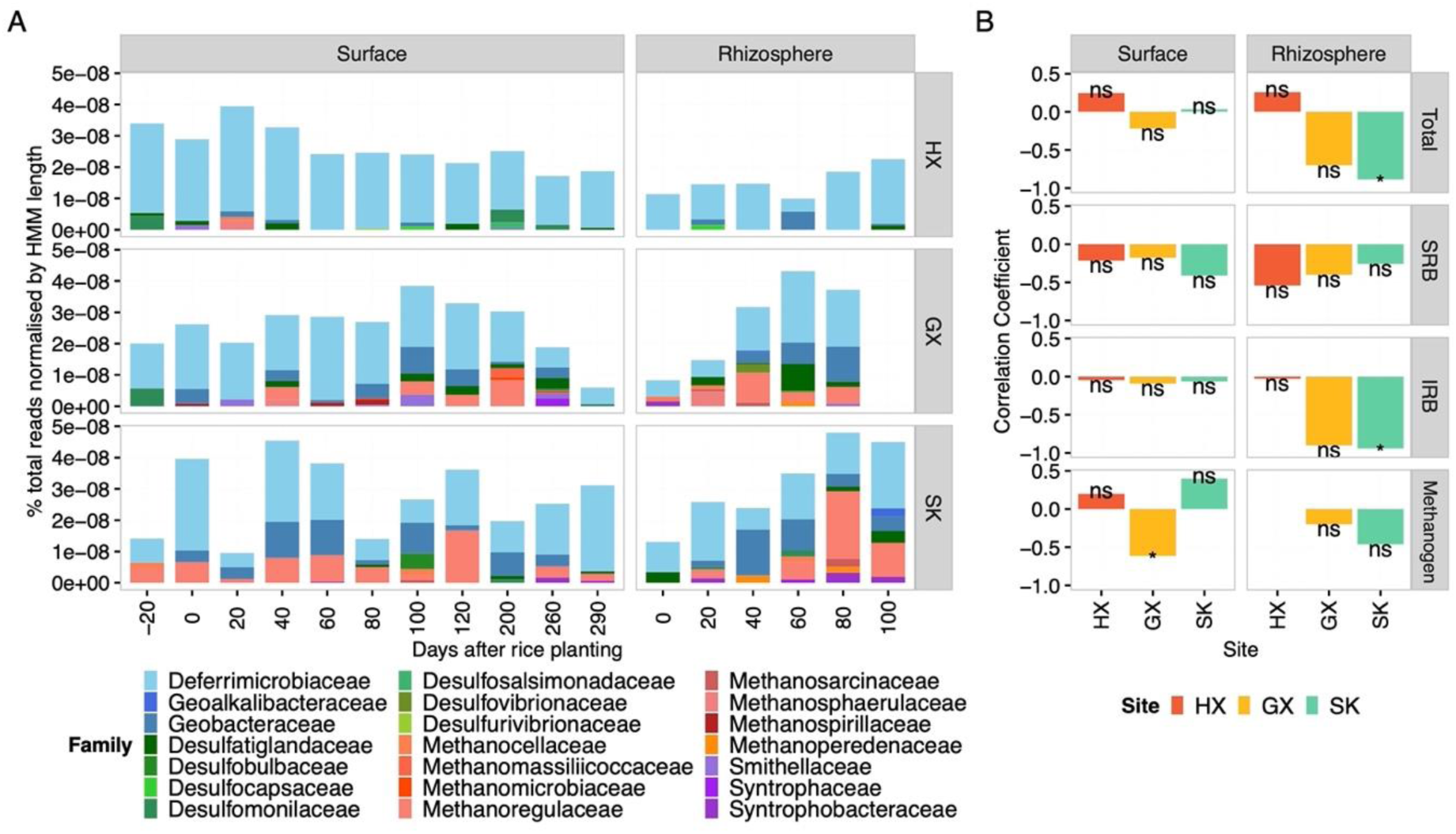
Analysis of *hgcA* genes and their correlation with MeHg. (A) Taxonomic distribution and abundance of *hgcA* genes across specific metabolic guilds throughout the sampling period. Color patterns represent iron-reducing bacteria (IRB, blue patterns), sulfate-reducing bacteria (SRB, green patterns), methanogens (orange/red patterns), and syntrophs (purple patterns). Note that sequences not classified to known functional groups were excluded from panel A for visualization purposes but are included in total abundance calculations. (B) Spearman correlation coefficients between MeHg concentrations and gene abundances for different metabolic guilds. Syntrophs are not shown due to insufficient data. Asterisks indicate statistical significance levels (* p<0.05; ns: not significant).

Temporally, *hgcA* abundance at HX remained relatively stable, mirroring MeHg trends (Fig. 1B). Across the growing season, correlations between total *hgcA* abundance and soil MeHg were inconsistent with guild-specific differences (Fig. 2B). Taxonomically, *hgcA* was primarily assigned to *Deferrimicrobiaceae* at HX, whereas *Methanoregulaceae*-related *hgcA* increased at GX and SK, particularly in rhizosphere samples.

As an additional test of drivers of MeHg accumulation, we fitted multiple linear regression models with model selection by AICc (Supplementary Result 2). The best-supported model included additive effects of bioavailable Hg(II) (F1-Hg) and overall *hgcA*, while controlling for site and soil compartment. In this model, F1-Hg was a major positive predictor of MeHg concentrations (β=0.73, p<0.001), whereas total *hgcA* abundance was not significant. Adding an interaction between F1-Hg and *hgcA* did not improve fit (ΔAICc=3). Consistent with this, the explained variance (adjusted R^2^) increased from ∼60% (additive model without site/compartment) to ∼84% when site and compartment effects were included, underscoring other environmental influences on MeHg.

These results suggest that net MeHg accumulation is not primarily controlled by methylation potential in Hg-contaminated rice paddies, but rather by Hg substrate availability and other environmental factors. This contrasts with findings from river systems where direct correlations between *hgcA* abundance and MeHg production have been observed under more uniform geochemical conditions (68–70). In rice paddies, the relationship between *hgcA* and MeHg content is complicated by several factors: (i) Pronounced geochemical gradients in DOM characteristics, redox conditions, and Hg bioavailability; (ii) the presence of multiple MeHg sinks including biotic and abiotic demethylation processes (20,71); and (iii) MeHg uptake by rice plants, which demonstrate efficient MeHg bioaccumulation from paddy soil (i.e., bioaccumulation factors (BAFs) of 5.5-5.6) (72,73). Although the quantitative effect of plant uptake on soil MeHg concentrations was not considered in this study, plant-mediated removal may partially contribute to soil MeHg content we observed. The interplay between MeHg production, degradation, and plant uptake processes operating simultaneously in rice paddies underscores the need to consider the genetic potential of methylating microbes alongside broader ecosystem variables that operate at larger spatial scales to better predict the fate of MeHg in rice paddy systems.

### *merB*-mediated demethylation and MeHg dynamics

Following *hgcA* analyses, we investigated demethylation potential represented by *merB*, which encodes the organomercurial lyase that performs reductive demethylation (RD) (74,75). Total *merB* gene abundance in rhizosphere was consistently higher compared to surface soil, which were similar across the three sites (HX:mean=5.1×10⁻⁸±5.5×10⁻⁸, n=11; GX: 3.5×10⁻⁸±5.2×10⁻⁸, n=11; SK:5.8×10⁻⁸±3.9×10⁻⁸, n=11) (Fig. S5). The consistently higher *merB* abundance in rhizosphere samples is likely attributed to radial oxygen loss (ROL) (76), creating a more oxidizing condition conducive to aerobic *merB-*carrying microbes (77), although direct soil ORP measurements were not obtained in this study. The highest rhizosphere *merB* was observed at day 0, followed by a gradual decline, aligning with previous findings that ROL rates decrease over the rice growth period (77). This temporal trend further supports the potential role of plant-mediated oxygen dynamics in structuring the *merB*-carrying community

Multiple regression analysis revealed that *merB* gene abundance has a significant positive association with MeHg levels (β=2.51×10^6^, p=0.0113) after accounting for site-specific effects. The positive association between gene abundance and absolute MeHg concentrations suggest adaptation of microbial communities to Hg-contaminated environments. In highly polluted aquatic systems, increased THg load drives higher absolute MeHg concentrations (74). The microbial communities respond by enriching for Hg-resistant bacteria carrying the *mer*-operon, including *merB* genes (74).

Crucially, this correlation between *merB* and MeHg is particularly strong in GX (Spearman’s ρ=0.6; n=5) and SK (Spearman’s ρ=0.83; n=6) rhizosphere samples. This suggests that the ROL-driven oxidizing conditions select for the aerobic *merB*-carrying organisms, and where THg loads are high (GX & SK), these adapted populations are most enriched, leading to the strong co-variance between *merB* gene abundance and MeHg concentration in this specific micro-environment.

At the transcriptomic level, *merB* transcripts were detected only in HX surface soils (n=6, Fig. S4B). This disconnect from gene abundance likely reflects: (i) site-and compartment-specific conditions regulating *merB* induction (e.g., oxygen and DOM), which may have been favorable only at HX surface at the sampling time (78); (ii) the transient nature of *merB* transcription and the short half-lives of *mer* transcripts, making them easy to miss in single time-point sampling; and (iii) temporal fluctuations in Hg(II) bioavailability that narrow the window for detectable *merB* expression (74,79). These findings highlight the need for temporal transcriptomic sampling to better establish relationships between *merB* expression and MeHg dynamics *in situ*.

### Methane-cycling communities show coupled dynamics and shared responses to Hg contamination

Recently, the role of methane-cycling microbes in mediating MeHg dynamics has become increasingly evident through controlled laboratory studies (14,22,80–82) However, the environmental relevance and field-scale implications of these laboratory-derived mechanisms remain largely unexplored in complex agricultural systems.

We show that despite contrasting Hg contamination sources (GX: atmospheric deposition; SK: historical smelting residues), methane-cycling communities at GX and SK exhibited similar beta-diversity (PERMANOVA, p<0.001; Fig. 3C&D). Constrained ordinations showed that geochemical variables explained 27.67% of the beta-diversity in methanogens and 22.14% in methanotrophs (p=0.001 and 0.004, respectively; Fig. 3C–D).

**Figure 3.**
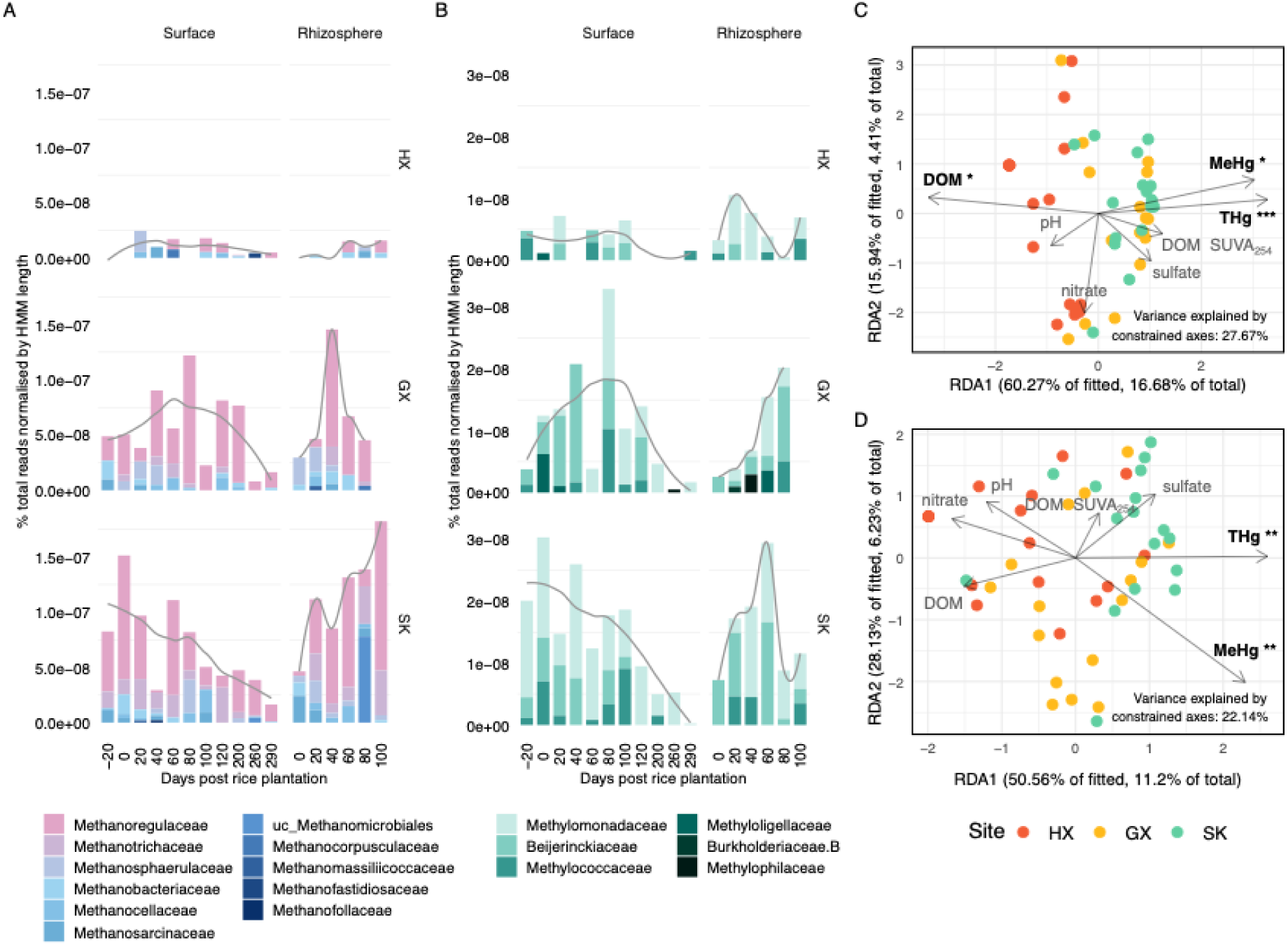
Community structure and environmental drivers of methanogens and aerobic methanotrophs. (A) Taxonomic distribution and relative abundance of *mcrA* genes across sampling days. (B) Taxonomic distribution and relative abundance of *pmoA*/*mmoX* genes across sampling days. (C&D): Distance-based redundancy analysis (db-RDA) of beta-diversity in relation to geochemical variables, with panel C showing methanogens and panel D showing methanotrophs. Points represent individual samples colored by site. Arrows indicate environmental variables with significance levels (*** p<0.001, ** p<0.01, * p<0.05).

THg and MeHg were significant predictors for both groups, and DOM concentration showed an additional significant association with methanogen beta-diversity (permutation tests, p<0.05; Supplementary Result 3).

Methanogen and methanotroph abundances were significantly and positively correlated (β=0.51, p=0.003, n=50; Fig. S6), indicating that their populations tended to co-vary across samples. However, methanotrophs were consistently present at roughly an order of magnitude lower abundance than methanogens (Fig. 3A&B). This observation aligns with studies demonstrating positive co-occurrence networks between these groups (83) and synchronized responses to environmental factors in rice paddies (84), likely reflecting their metabolic interdependence. Methanotrophs rely on methane produced by methanogens and may, in turn, help regulate local oxygen concentrations (83,84). This interdependence is further supported by the notably lower abundances of both groups at HX, the site with the highest average ORP, where conditions appeared too oxidizing for robust methanogenesis and suboptimal for aerobic methanotrophy, resulting in reduced populations of both metabolically linked functional groups.

The similarity of methane-cycling community structures at Hg-impact sites, in addition to their significant association with THg and MeHg suggests the presence of a core methane-cycling community that adapts similarly to Hg impacts regardless of contamination source.

### Context-dependent methanogen-MeHg relationships links to DOM quality and compartment effects

We subsequently examined how methanogen abundance (via *mcrA*) links to MeHg concentrations across sites and soil compartments. Overall methanogen abundance showed marginal association with MeHg (β=-1.30×10^7^, p=0.064), but this relationship varied with THg concentration (β=4.57×10^2^, p=0.049), indicating context-dependent relationships where higher methanogen abundance amplifies MeHg production under elevated THg conditions.

Methanogen community beta-diversity correlated significantly with DOM (Fig. 3C). Additionally, methanogen abundance showed negative correlations with both DOM quantity (β=-5.94×10^-8^, p=0.007) and SUVA_254_ (β=-2.88×10^-8^, p=0.003), suggesting lower abundance where DOM is less bioavailable. Similarly, DOM SUVA_254_ correlated negatively with MeHg (β=-0.48, p=0.005) while DOM quantity showed no significant relationship (β=0.60, p=0.140), aligning with previous work showing that DOM bioavailability more critically determines MeHg concentrations than quantity (10,67).

Our correlational field data reveal significant associations between DOM, methanogens, and MeHg, these observational relationships that are consistent with established experimental evidence for DOM’s dual role in methylation processes. Higher SUVA_254_ values indicate aromatic, humified DOM that reduces MeHg through strong Hg complexation (10,67). Conversely, bioavailable DOM fractions (autochthonous DOM, fresh organic matter, low-molecular-weight acids, algal organic matter) enhance MeHg production by providing carbon sources and stimulating methylating microorganisms, including methanogens (10,63,67). The negative correlations with SUVA_254_ support this framework: Highly aromatic DOM suppresses both methanogenic proliferation and MeHg formation, while bioavailable DOM promote both processes.

Specific methanogen taxa showed compartment-dependent patterns. *Methanoregulaceae*, the dominant family across sites with diverse Hg methylators (7), (Fig. 3A, Supplementary Result 4), mirrored overall methanogen trends with positive MeHg correlations in SK surface soil (ρ=0.745, p=0.011; n=11) but negative correlations in SK rhizosphere (ρ=-0.943, p=0.017; n=6). These family-specific *mcrA* patterns differ from the relationships shown in Fig. 2, which reports methanogen-associated *hgcA*; methanogens may directly or indirectly contribute to methylation and demethylation, so *mcrA*–MeHg associations do not necessarily mirror *hgcA*–MeHg patterns.

Minor taxa showed different compartment preferences. *Methanospirillaceae* (ρ=0.974, p=0.005; n=6) and *Methanosarcinaceae* (ρ=0.941, p=0.005; n=6) correlated positively with MeHg in SK rhizosphere, with weaker patterns at HX and GX. Metatranscriptomic analysis (n=6) detected active *mcrA* expression across sites, with higher levels at Hg-contaminated sites, particularly by *Methanoregulaceae*, providing functional support for gene abundance data (Fig. S4C).

Interestingly, we recovered sequences from *Methanoperedenaceae*, an archaeal group (ANME), which performs anaerobic oxidation of methane (AOM) via reverse methanogenesis (85). This finding corroborates the detection of *Methanoperedenaceae*-associated *hgcA* from our rice paddy metagenomes (Fig. 2A, Supplementary Result 7). Although *mcrA* and *hgcA* transcripts associated with *Methanoperedenaceae* were detected at the Hg-impacted sites, (Fig. S4C), we exclude this group from subsequent analyses due to the lack of empirical evidence for their methylation capacity. Their potential role in Hg cycling are discussed in Supplementary Discussion 1.

### Flooding associates higher methanotroph abundance with lower MeHg

Methanotroph abundance relationships with MeHg followed similar site-specific patterns to those observed for methanogens, consistent with the two guilds co-varying across samples (Fig. A&B). However, when focusing specifically on the flooded period, multiple regression revealed a significant negative association between methanotroph abundance and MeHg concentrations (β=–3.21 × 10⁷, p=0.009), after adjusting for site and soil compartment, an effect not observed in the methanogen data.

Metagenomic analyses revealed that *Methylomonadaceae*, *Beijerinckiaceae*, and *Methylococcaceae* were the most abundant methanotroph taxa (Fig. 3B, Supplementary Result 5). Metatranscriptomic data further confirmed active methanotrophy at all sites, with particularly high *pmoA* expression at SK, where *Methylomonadaceae* was the most highly expressed methanotroph family (Fig. S4D). *Methylomonadaceae* showed the strongest negative correlations with MeHg at GX (ρ=-0.8, p=0.133; n=5) and SK (ρ=-0.9, p=0.083; n=5) rhizosphere, while other methanotroph taxa showed weaker correlations (Supplementary Result 6).

It is likely that the negative methanotroph–MeHg relationship reflects either (i) flooding-induced oxygen depletion that suppresses aerobic methanotrophs while stimulating anaerobic Hg methylation, or (ii) the influence of taxa such as *Methylomonadaceae*, which are adapted to low-oxygen conditions via anaerobic denitrification (86–88), allowing persistence in flooded, microaerophilic rhizospheres and potential methanotroph-mediated demethylation via a methanobactin-related pathway, which we explore in the next section.

### Exploratory analysis of *mbnT* as a potential demethylation biomarker

Recent studies indicate a methanotroph-specific MeHg demethylation pathway facilitated by methanobactins (24,25). Methanobactins are copper-sequestering chalkophores essential for particulate methane monooxygenase (pMMO) activity in some methanotrophs (89). While primarily binding copper with high affinity, methanobactins also bind other metals, including Hg(II), albeit with lower affinity (89).

The cellular uptake of methanobactins is mediated by MbnT, a TonB-dependent transporter (TBDT), which is an outer membrane protein (90). This uptake is crucial as methanobactin appears to deliver MeHg inside the cell, and a study shows that deleting *mbnT* impairs methanobactin uptake and, consequently, MeHg degradation (24,25). The degradation of MeHg is thought to be carried out by the periplasmic methanol dehydrogenase, targeting the carbon-Hg bond, and is distinct from the *merB*-dependent organomercurial lyase pathway, as these methanotrophs generally lack the *merB* gene (24,25)

Given the prevalence of methanotrophs in rice paddies, we investigated whether their observed association with MeHg dynamics is linked to MbnT-mediated demethylation. We examined the genomic context of previously studied demethylating methanotrophs and methanotroph-MAGs recovered from rice paddies to determine if they encode genes related to methanobactin biosynthesis (*i.e., mbnA, mbnB, mbnC* as core components, and additional enzymes like *mbnD, mbnF, mbnN, mbnS, mbnX*) and transport (*mbnE, mbnM, mbnT*) (47). Aligning with previous research, we confirmed that the demethylating *Methylosinus trichosporium OB3b*, encodes the full suite of genes for methanobactin biosynthesis and uptake/transport (Fig. 4). *Methylomicrobium album BG8* and *Methylocystis sp. strain Rockwell*, also able to demethylate, encode *mbnT* but lack genes necessary for methanobactin biosynthesis. Conversely, *Methylococcus capsulatus Bath* lacks genes for both methanobactin biosynthesis and uptake (*mbnT*) and was shown to not demethylate (Fig. 4) (24). Subsequently, we show that none of the methanotroph MAGs possessed *mbnT* genes sufficiently similar to those found in the experimentally confirmed demethylating methanotrophs, and these MAGs also lacked all essential genes for methanobactin biosynthesis. Therefore, the rice paddy MAGs are unlikely to exhibit MeHg demethylation activity, although some of these MAGs showed significant correlations with MeHg via network analyses (Supplementary Result 8 & Discussion 2).

**Figure 4.**
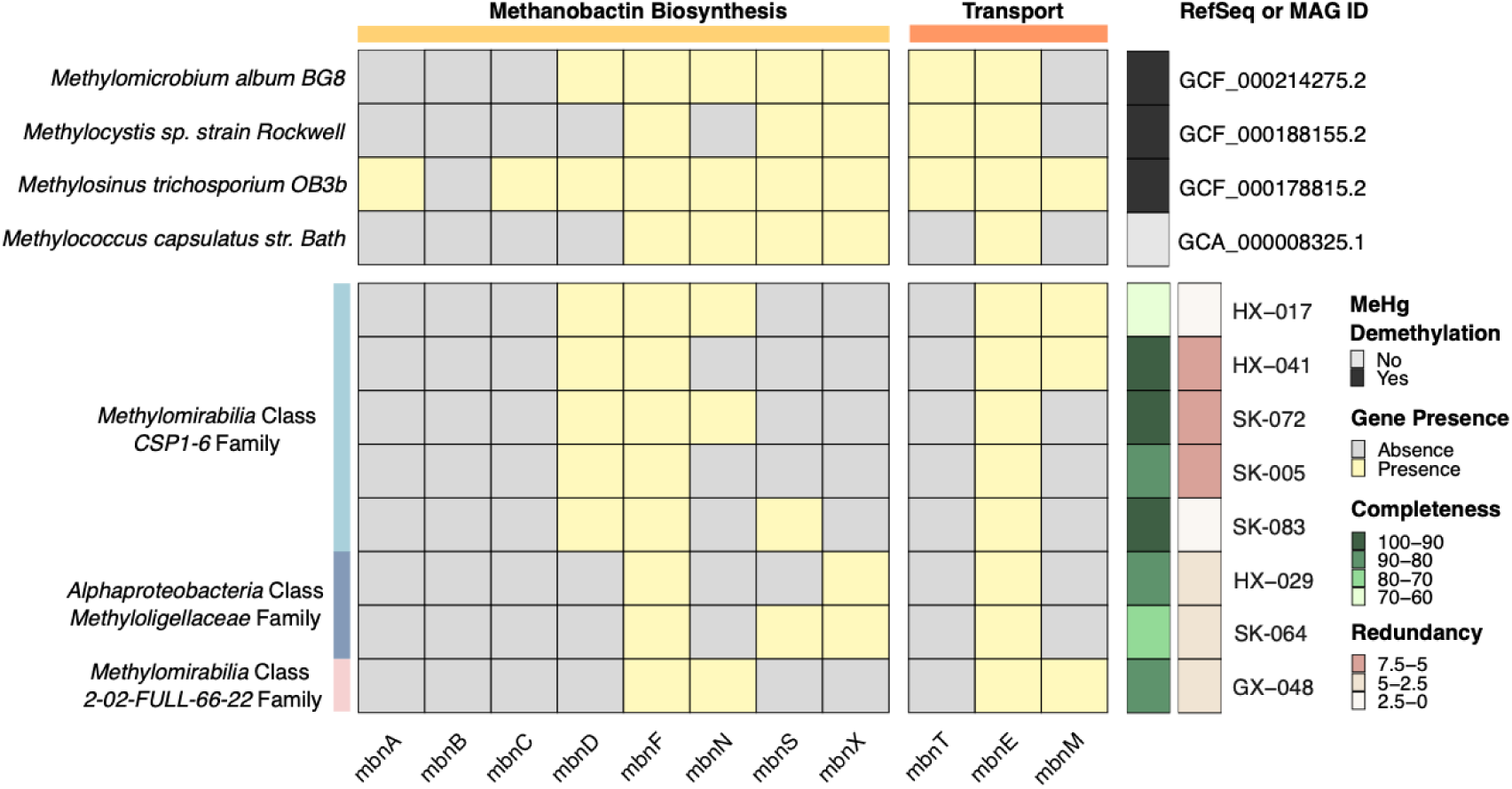
Presence/Absence of Methanobactin-Related Genes in Methanotroph Genomes. This heatmap illustrates the presence and absence of methanobactin biosynthesis and transport genes across experimentally studied methanotrophs (24) and MAGs recovered in this study, along with their respective quality metrics. The x-axis denotes the methanobactin genes, while the left y-axis represents the taxonomic classification of each genome, and the right y-axis lists the RefSeq or MAG IDs. All RefSeq genomes lack MBn biosynthesis, except OB3B (Note *mbnB* is present is OB3B as a pseudogene, which is not recovered through the HMM based analyses, but confirmed with RefSeq annotation).

Based on experimental evidence, we hypothesized that *mbnT* could serve as a marker for MeHg demethylation capacity as the demethylating methanotrophs are shown to possess the gene and subsequently analyzed *mbnT* from assembled metagenomes of rice paddies. We recovered *mbnT* that resemble those found in known demethylating methanotrophs using an HMM (TIGR01783, sequence cutoff 223.5). However, recognizing that TBDTs, including *mbnT*, have homologs in diverse organisms beyond methanotrophs, we employed a conservative approach, retrieving sequences classified to known demethylating methanotroph genera: *Methylomicrobium*, *Methylocystis*, and *Methylosinus* (Fig. S7). Transcriptomic analyses show that *mbnT* transcripts from these genera were present in Hg-contaminated rice paddies, though to a limited extent (n=6, Fig. S4E). Multiple regression analyses revealed that total *mbnT* gene abundance from these genera represents a significant positive predictor of MeHg concentration (β=4.579 ×10⁷, p=0.0136) after accounting for site and compartment specific effects (Supplementary Result 9), contrary to an expected negative correlation if *mbnT* represents effective demethylation.

Our findings suggest that while methanotroph demethylation capacity appears to be present in rice paddies, utilizing *mbnT* as a marker to assess net MeHg accumulation requires consideration of several limitations: (i) *mbnT* is not single-copy gene (91), and it belongs to the family of TBDTs, which are responsible for the uptake of diverse substrates, including metal chelates, and have numerous homologs in organisms other than methanotrophs (47).

Despite using stringent cutoff, the presence of *mbnT* homologs with high similarity could lead to biased abundance estimations. Furthermore, gene abundance dose not directly equate to activity. (ii) Methanotrophs exhibit varied demethylation rates (24). These rates depend on various biological factors, including the structure of methanobactin (24), the efficiency of the MbnT transport system (89), and the presence of additional copper-handling proteins, like MbnH and MbnP (90), as well as environmental factors such as redox conditions, pH, and the presence of competing metals (92–94). (iii) Methanobactins can influence Hg cycling through secondary mechanisms. This includes altering microbial community composition (*e.g.,* through copper competition impacting denitrifying bacteria (89)) or affecting the bioavailability of Hg species via competitive binding (94,95). These broader effects complicate the interpretation of *mbnT’*s role in MeHg demethylation.

Future studies should prioritize establishing direct relationships between *mbnT* expression and demethylation rates *in situ* through quantitative approaches with adequate sample sizes, before linking *mbnT* abundance to environmental MeHg dynamics. Additionally, broader experimental validation across diverse methanotrophic taxa is needed to confirm the universal applicability of this biomarker. Given the complexity revealed by our environmental data, controlled laboratory studies examining *mbnT* expression under varying Hg, redox, and competing metal conditions would help define when *mbnT* serves as a reliable indicator of methanotroph-mediated MeHg demethylation.

### Methane-cycling microbes as key biological predictors of MeHg variability

Our analyses demonstrate that relying on single factors has limited explanatory power for net MeHg accumulation, despite significant associations found for individual variables. The complexity expected from concurrent methylation and demethylation processes is reflected in the increase in model R² when accounting for site and compartment effects (e.g., from 60% to 84% in hgcA models). To address the limitations of single-factor analyses and identify the key, potentially non-linear, variables driving MeHg patterns across diverse functional genes and varying environmental conditions, we employed Random Forest (RF) analysis.

The RF models built using all variables achieved moderate predictive performance (R²=0.70±0.13, range: 0.40-0.95, Fig. 5). Bioavailable Hg(II) (F1-Hg) consistently emerged as the dominant predictor across all data splits (100% significance rate, p=0.010). Methanogen abundance ranked as the second most important variable (Mean IncMSE=20.8, 100% significance rate, p=0.013), followed by methanogen-hgcA abundance (Mean IncMSE=9.3, 49% significance rate), substantially outperforming other microbial indicators like total Hg methylators (*hgcA*), sulfate-reducing bacteria (SRB), and iron-reducing bacteria (IRB) abundances.

**Figure 5.**
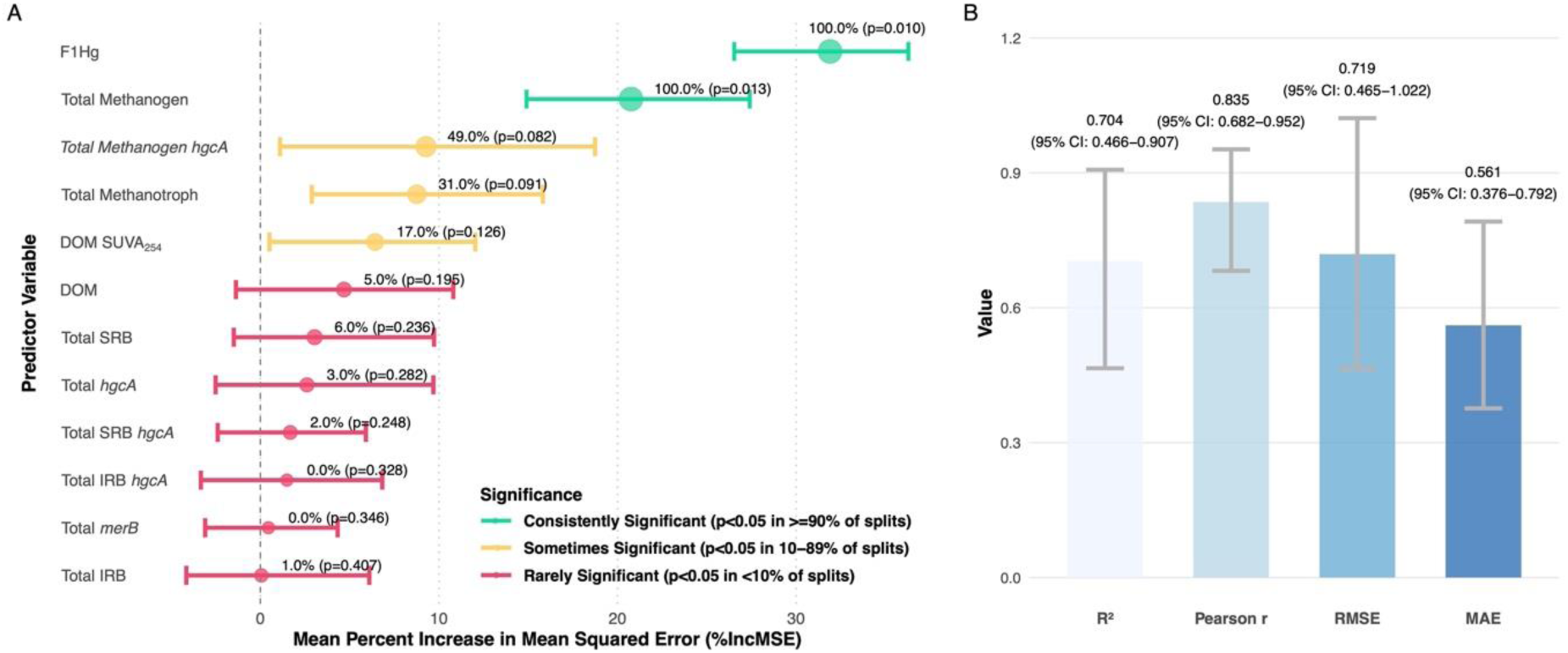
Variable importance and model performance for predicting MeHg concentrations using random forest analysis (full model). (A) Variable importance plot showing the mean percent increase in mean squared error (%IncMSE) for each predictor variable when permuted, with 95% confidence intervals across 100 different data splits. Larger point size and green coloration indicate higher significance rates. Values show the percentage of data splits where the variable was significant (p<0.05) followed by the mean p-value. (B) Model performance metrics displaying mean values with 95% confidence intervals across all data splits. The model was built using 2001 trees with mtry=4 and validated through 100 permutation tests with random 70/30 train/test splits.

Methanotroph abundance was identified as the third most important biological variable overall (Mean IncMSE = 8.8; 31% significance rate). Crucially, an explicit interaction term between methanogen and methanotroph abundances (i.e., their product) was a critical predictor of MeHg (Mean IncMSE = 18.3; 80% significance rate; p=0.038). This result suggests that the methanotrophic influence is primarily expressed through community-level coupling with methane-producing populations. Given that methanogens are key methylators that amplify MeHg under elevated THg, and methanotrophs showed a significant negative association with MeHg during the flooded period (suggesting a role in MeHg consumption/demethylation), the interaction term highlights that the net MeHg outcome is highly sensitive to the balance and synchronous activity of these two metabolically interdependent microbial guilds.

Constraining the RF model solely to the flooded period substantially improved predictive accuracy (R^2^=0.86±0.08; Fig. S8B). This improved performance was accompanied by a modest increase in the relative importance of methanogens (mean %IncMSE rose from approximately 21 to 23) and a decrease for F1-Hg (mean %IncMSE fell from approximately 32 to 26), suggesting that the predictive power of methanogens is comparatively greater under flooded, reduced conditions than that of Hg bioavailability alone. Integrating genomic and geochemical data through RF modeling revealed that methanogen, methanogen methylator, and methanotroph abundance were the top biological factors contributing to net MeHg accumulation, while Hg bioavailability and dissolved organic matter (DOM) aromaticity were the dominant geochemical factors. Prediction accuracy further improved when removing less influential variables, reinforcing the importance of these key factors.

While previous studies have associated methanogenesis with MeHg content in rice paddies (14,21), our work advances this understanding by identifying methanotrophy as an important biological correlate of net MeHg accumulation and characterizing specific methane-cycling populations associated with MeHg variability. Methanotrophic involvement is particularly noteworthy, as it suggests a previously underappreciated potential demethylation pathway in rice paddies. Our findings indicate that studying the relationship between microbial methane metabolism and Hg cycling provides critical insights into the key biogeochemical factors associated with net MeHg accumulation in Hg-contaminated rice paddies.

While these findings advance our understanding of microbial associations with MeHg dynamics, several considerations should guide their interpretation and application. The observed coupling between methane and Hg cycling, though mechanistically plausible, requires validation across diverse environmental contexts before broader generalization to other rice paddy systems. Our RF approach, while effective at identifying statistical associations, cannot establish causal directionality between predictors and MeHg accumulation, highlighting the need for controlled experimental validation of these relationships. We recommend validation across diverse rice paddy systems with (i) varying flooding patterns and water management practices (12,96), (ii) different rice cultivars (97), and (iii) diverse geographical span (3) and contamination sources (98–100) to benchmark model transferability.

Our results underscore that effective assessment of MeHg risk requires integrating microbial community data with physicochemical gradients that govern both Hg bioavailability and microbial niche occupation. The importance of methane-cycling communities in our predictive models highlights promising avenues for management strategies that could simultaneously address climate change mitigation and Hg pollution through targeted manipulation of shared microbial pathways (101).

## Acknowledgement

This research was funded by the National Natural Science Foundation of China (Grant No. 42394092), the Guizhou Provincial Major Scientific and Technological Program ([2024]013) to BM, and the Natural Sciences and Engineering Research Council of Canada to AJP and DSG. We thank Tianli Zhan for contributing to the geochemical analyses. The bioinformatics analyses were conducted using computing clusters at the Digital Research Alliance of Canada.

